# Cell-type-specific characterization of miRNA gene dynamics in immune cell subpopulations during aging and atherosclerosis disease development at single-cell resolution

**DOI:** 10.1101/2023.10.09.561173

**Authors:** Ana Hernández de Sande, Tanja Turunen, Maria Bouvy-Liivrand, Tiit Örd, Senthil Palani, Celia Tundidor-Centeno, Heidi Liljenbäck, Jenni Virta, Olli-Pekka Smålander, Lasse Sinkkonen, Thomas Sauter, Anne Roivainen, Tapio Lönnberg, Minna U Kaikkonen, Merja Heinäniemi

## Abstract

MicroRNAs (miRNAs) are a class of regulatory non-coding RNAs that finetune cellular functions by modulating the stability and abundance of their target mRNAs, thereby contributing to regulation of tissue homeostasis. MiRNA genes are transcribed similarly to protein-coding genes and recent studies have enabled their annotation and quantification genome-wide from bulk nascent transcriptomes. Here, we developed an approach to quantify and integrate miRNA gene signatures into single-cell studies. To characterize miRNA gene expression dynamics, we first compared the suitability of droplet and plate-based single-cell RNA-sequencing (scRNA-seq) platforms using the matched datasets provided by the Tabula Muris Senis and Tabula Sapiens consortiums. We found high concordance between the platforms and with cell type-specific bulk expression data. Based on the comprehensive aging profiles, our analysis comparing spleen immune cells between young and old mice revealed a concordant regulation of miRNAs involved in senescence and inflammatory pathways in multiple immune cell types, including up-regulation of mmu-mir-146a, mmu-mir-101a and mmu-mir-30 family genes. To study the aberrant regulation of immune cell homeostasis and tissue inflammation that pre-dispose to aging-related disease development, we collected transcriptome profiles from atherosclerosis development in LDLR^-/-^ApoB^100/100^ mice. We found an elevated myeloid cell proportion in the adipose tissue and further characterized the cell subtypes based on reproducible transcriptome clusters. We then compared miRNA gene expression in early versus late disease and upon inflammatory challenge to monitor different stages during disease progression. At atherosclerotic stage, pro-inflammatory mmu-mir-511 expression increased in several macrophage subtypes, while immunosuppressive mmu-mir-23b∼mir-24-2∼mir-27b up-regulation was specific to Trem2+ lipid-associated macrophages. The infiltrating monocytes up-regulated mmu-mir-1938 and mmu-mir-22 expression and in classical monocytes maturation further increased mmu-mir-221∼222, mmu-mir-511 and mmu-mir-155 expression. To validate that these changes detected from single cell profiles represent miRNA gene transcriptional regulation, we used nascent transcriptomics data from *ex vivo* macrophage cultures with pro-inflammatory stimulation, confirming both rapid and long-lasting transcriptional activation of the miRNA loci studied. Collectively, our work enables integrating miRNA gene analysis to current single cell genomics pipelines and facilitates characterization of miRNA regulatory networks during aging and disease development.

## INTRODUCTION

The study of single-cell (sc) transcriptomes has revolutionized the field of cell biology, enabling identification of new cell types, cellular states and characterizing cellular transitions across healthy tissues and during disease development ^1^. MicroRNAs (miRNAs), a class of regulatory non-coding RNA molecules, can base pair to their target messenger RNA (mRNA), thereby interfering with their translation into proteins. Thus, miRNA-mediated post-transcriptional regulation strongly impacts gene-regulatory networks that modulate cell function via controlling cell homeostasis ^2^. At systems level, control of cell state transitions deteriorates over time, impairing cellular homeostasis and tissue function. This process manifests with low-grade tissue inflammation and constitutes a risk for developing inflammatory-related diseases such as diabetes, atherosclerosis, Alzheimer’s disease, and certain cancers (reviewed in ^3,4^).

MiRNA genes correspond to long transcripts called primary miRNAs (pri-miRNA) that are transcribed by RNA polymerase II similarly as protein-coding genes. Subsequently, miRNA transcripts are processed into short transcripts, pre-miRNAs, and further into 20-22 nucleotides (nt) long mature miRNAs. During sample preparation for standard bulk RNA-sequencing, the size selection step excludes the processed miRNA transcripts. Therefore, separate protocols for small RNA sequencing have been developed and represent the most common miRNA profiling method (reviewed in ^5^). Recently, these were adapted to achieve cellular resolution, however the feasibility of sc-small RNA-seq is limited by low throughput ^6,7^.

Presently, the comprehensive gene annotations such as Refseq primarily consist of pre-miRNA coordinates, appropriate for small RNA sequencing data analysis rather than conventional transcriptome sequencing. As the reference annotation is commonly utilized in single-cell studies, quantification of miRNA genes and the analysis of their regulation at cell type resolution is lacking. In consequence, our understanding of how miRNAs integrate into regulatory networks that govern cell state homeostasis is incomplete. Acknowledging that single-cell transcriptomics captures between 17 to 23% of unspliced reads ^8^, analysis of pri-miRNA transcripts presents an alternative. In our previous work, we developed a comprehensive miRNA gene annotation approach based on nascent transcriptome (Global-run-on coupled with sequencing, GRO-seq), Cap Analysis of Gene Expression (CAGE) and histone marker data that enabled the quantification of pri-miRNA transcriptional activity in a multitude of bulk genomics studies in cell lines and primary tissue contexts ^9,10^.

Here, we leverage our previous approach annotating miRNA gene coordinates to quantify miRNA genes from single-cell transcriptomes. To discover changes in miRNA gene expression that could impact their central role in control of cellular homeostasis, and thereby contribute to the progressive loss of healthy physiology, we used the comprehensive aging mouse dataset, collected by Tabula Muris Senis (TMS) consortium ^11–13^, and novel profiles from an atherosclerosis disease model. We demonstrate how miRNA gene activity is impacted in immune cells by aging and during disease development and provide these datasets and annotations as an openly available resource to facilitate further characterization of miRNA regulatory networks.

## RESULTS

### Quantification of miRNA gene expression in single-cell transcriptomes

To characterize miRNA gene expression dynamics in single cell transcriptomes collected from mouse tissues, we first followed the approach described in ^9^ to obtain mouse transcript coordinates representing intergenic (transcribed from their own promoter) and intragenic miRNA genes that are co-transcribed from introns of their host genes (see Methods, datasets used are listed in Table S1A). For simplicity, miRNA genes transcribed from alternative transcription start site (TSS) were summarized by gene locus, following the current practice of capturing gene-level expression in single-cell datasets (see Methods, similarly human coordinates were adopted from ^9^). This annotation, which resulted in 233 intragenic and 135 intergenic miRNA gene loci (368), corresponding to 990 mature miRNAs for mouse, and 511 and 391 loci, respectively, for human (1896 mature miRNA), was integrated into the GenCode transcript annotations (Fig. 1A, Table S1B-C: miRNA gene coordinates).

**Figure 1.**
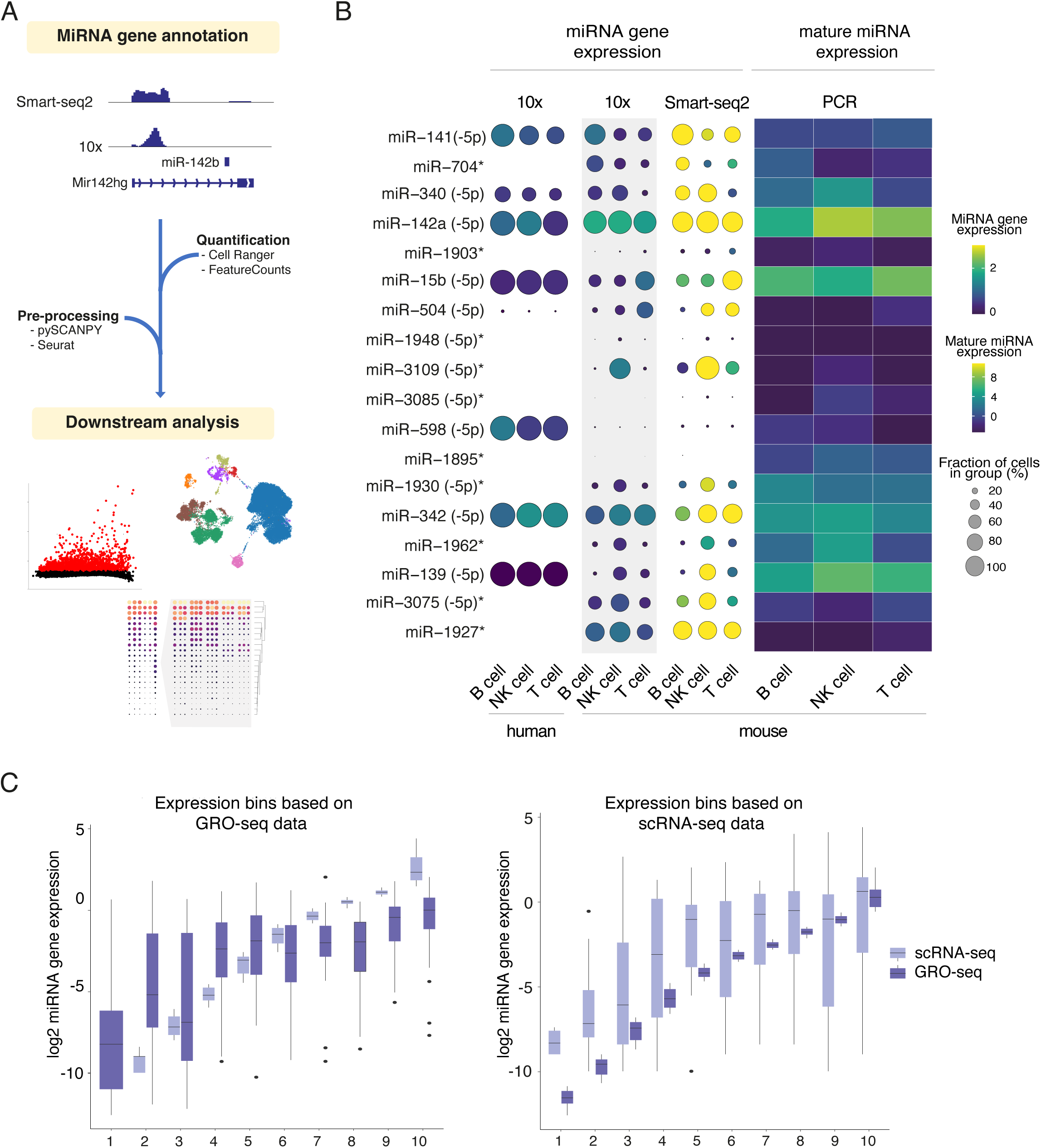
MiRNA gene annotation and quantification in scRNA-seq datasets. (A) Overview of the miRNA gene quantification for scRNA-seq data. Gene and miRNA gene counts were extracted with the Cell Ranger or FeatureCounts pipelines, followed by downstream analysis performed by combining Seurat ^88^ and SCANPY ^89^ packages to obtain and study cell-type-specific miRNA gene expression at single-cell resolution. (B) MiRNA gene markers (n=6 per cell type, defined from mouse droplet-based 10x Genomics data in grey shade) were compared to their corresponding human genes and mature forms in splenic cells including B-cell, NK-cell, and T-cell subpopulations. On the left, dot plot heatmaps shows miRNA gene expression (from high expression in yellow to low expression in blue) and percentage of expression (circle size) based on 10x Genomics and FACS-coupled Smart-seq2 technology in mouse. On the right, the heatmap depicts mature miRNA expression levels from FACS-sorted splenic cells measured with PCR (GSE144081 from ^16^). * mouse miRNA genes that were not annotated in the human genome. (C) Comparison of scRNA-seq-based quantification (log2 miRNA gene expression) to GRO-seq-based primary transcript expression in the mouse stromal cell line ST2. Genes were binned from low to high expression levels based on GRO-seq data (lower panel) or scRNA-seq data (upper panel).

Next, we retrieved the comprehensive aging mouse dataset, collected by the TMS consortium ^11–13^ to serve as the first benchmark for miRNA gene quantification at cellular resolution across tissues. Single-cell transcriptomes collected by TMS were sequenced with 10x Genomics and Switching Mechanism at the 5′ end of RNA Template (Smart)-seq2 technologies (Table S1C-D). While both methods follow a poly-A-based priming strategy, 10x Genomics uses 3’-based quantification whereas Smart-seq2 captures reads along the entire transcript ^14^. To assess which method could more accurately capture miRNA gene expression dynamics, we quantified read counts from miRNA gene coordinates in splenic cells from 3-month-old mice from both platforms (matching human data was retrieved from Tabula Sapiens ^15^). MiRNA gene dynamics were studied based on the ability of each platform (i) to measure miRNA gene expression levels, and (ii) the ability to detect miRNA quantified as ‘percentage of expression’.

In line with previous research, Smart-seq2 libraries performed worse on detecting different miRNA genes: across cell types, on average 225 miRNA were found with 10x Genomics vs. 66 with Smart-seq2, despite that more reads per miRNA gene detected were captured at the utilized sequencing depth (98 vs. 83 per miRNA detected, Fig. S1, ^7^). Overall, expression profiles across cell types correlated well between the two platforms, with a 0.83 correlation among miRNA gene average expression level genome-wide and 0.87 comparing the percentage of expression.

To study the relationship between miRNA genes and their corresponding mature forms in individual transcriptomes, we identified miRNA gene markers (n=6) for the main splenic immune cell populations including mouse B-, T-, and NK-cells using the statistical tests for cluster comparison in the Single-Cell Analysis in Python (SCANPY) pipeline (Fig. 1B, dot plot panels). We then retrieved their corresponding mature forms and quantified their expression from polymerase chain reaction (PCR)-based profiles available from the immune cell atlas (GSE144081 from ^16^; Fig. 1B, heatmap right panel). Relative expression levels amongst the platforms, across species and between the gene and corresponding mature forms were concordant for highly expressed genes, exemplified by (mmu)-miR-141 with highest expression in B-cells and mmu-miR-340 with high expression in NK-cells and B-cells in relation to T-cells (Fig. 1B, notice that miRNA loci marked * did not have corresponding human data). Overall, the correlations between the percent of expression or average expression in single-cell analysis and mature miRNA expression were generally weaker (0.44 - 0.47 in all the comparisons). However, as the mature miRNA levels are also affected by transcript processing and stability, we performed a more direct benchmark comparison to primary transcription assayed using bulk GRO-seq and parallel 10x Genomics scRNA-seq in the mouse stromal cell line ST2 (see Methods, Fig. 1C). MiRNA genes were divided into 10 bins based on their expression in each sequencing technology (plotted from low to high values in Fig. 1C, light purple indicates 10x Genomics-based scRNA-seq and dark purple GRO-seq signal). Independently of the data type used to bin miRNA genes, the detected expression level was highly comparable at bins of high expression (bins from 6 to 10), whereas the bins corresponding to lower expression levels displayed higher variability. This observation agrees with limitations in efficient capture of low-expressed transcripts in 10x Genomics scRNA-seq datasets ^7^. Taken together, quantifying miRNA gene expression based on 10x Genomics and Smart-seq2 scRNA-seq platforms has high concordance with cell-specific bulk expression data. Since more miRNA genes were detected from 10x Genomics-based profiles, we continued with this technology in downstream analyses to capture miRNA transcription at cellular resolution.

### Aging profiles in splenic immune cells reveal coordinated and cell-type-specific changes in miRNA gene expression

During aging, the immune system deteriorates, manifesting in loss of homeostatic mechanisms controlling immune responses that can underlie chronic inflammation and thereby risk for developing various aging-related disease. To gain insights into miRNA expression profiles in aging immune cell subpopulations, we retrieved mouse samples from the TMS consortium, the largest resource of single-cell datasets to study aging in multiple tissues. We focused on the splenic male samples that covered the broadest range of time points. The annotation contained seven major splenic cell types that cluster together independently of the mice age (Fig. 2A and Fig. S2A). During aging, mature NK-cell, T-cells, and plasma cells proportions increased whereas macrophage, proerythroblast, T-cell, and NK-cell proportions decreased (Fig. 2B and Fig. S2B), in line with previous results ^17^. Next, we compared within each cell type young (1 and 3 months) and old mice (24 and 30 months) to track progressive, gradual changes in gene expression that are detectable only after sufficient time ^13^ and distinguished in each cell type changes in the expression level within cells expressing the transcript (DE category) or variations in the percentage of cells expressing a particular gene (DZ category) (see Methods, n=6996 cells). In total 131 DZ and 58 DE miRNA genes were identified across the different cell types, summarized in Fig. 2C (Table S2). Next, miRNA genes were ranked based on combined p-values (Fisher’s exact test, see Methods) to identify concordant changes amongst the five most abundant immune cell types, which revealed 187 and 42 significant (combined p-value < 0.05) miRNAs in DZ and DE categories, respectively. A subset of 94 top ranked miRNA with log2 fold change in detection rate >0.5 in two cell types are shown in Fig. 2D (refer to Fig. S2C showing their profile in female samples).

**Figure 2.**
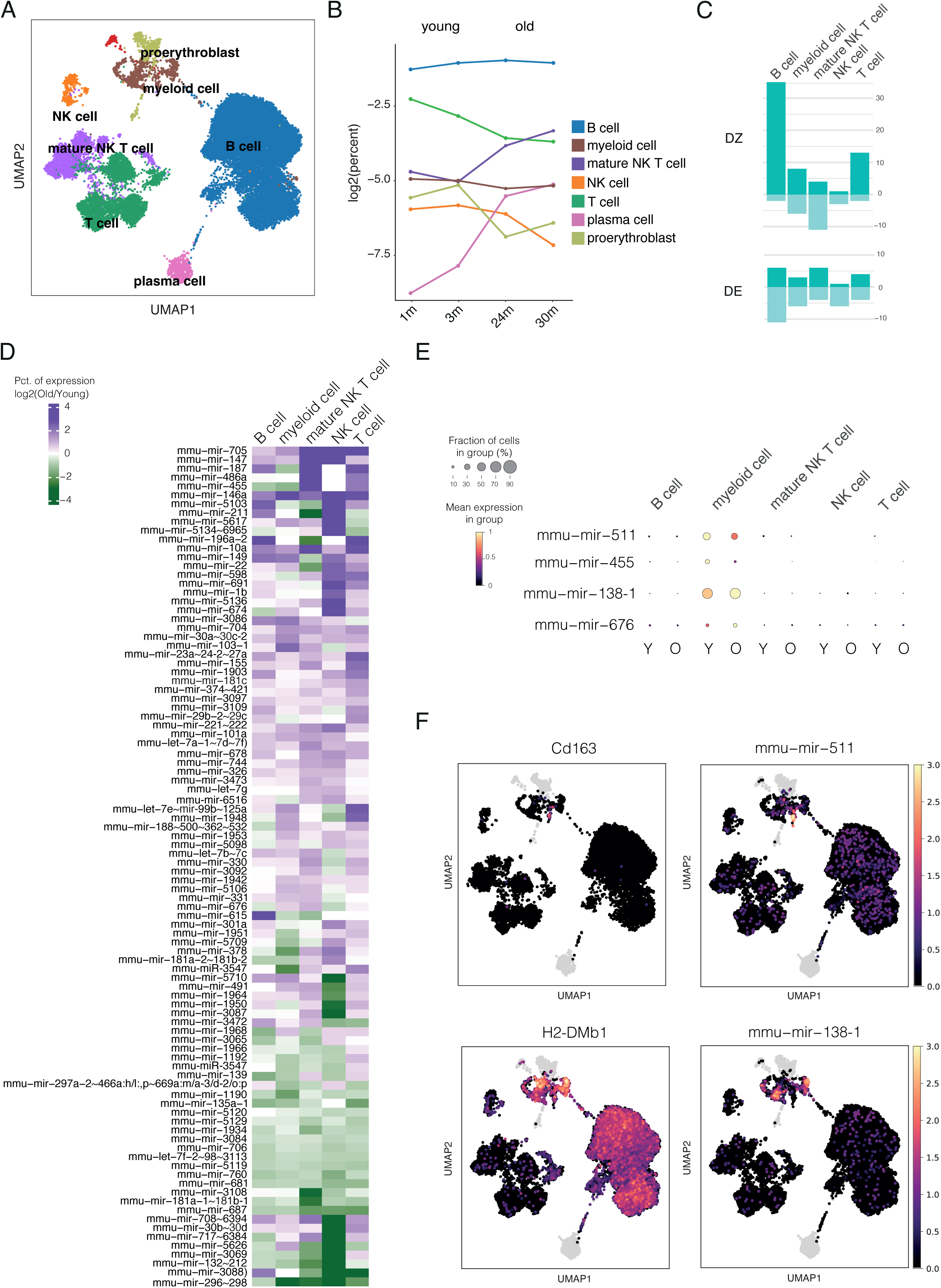
Differential expression analysis of miRNA genes in spleen tissue during aging. (A) Uniform manifold approximation and projection (UMAP) plot of the integrated spleen (all age groups) dataset (n=27260 cells). Colors indicating different cell types (annotations from TMS) from 8 major clusters (n>150 cells) include B cells (n=18398) in dark blue, T cells (n=4029) in forest green, mature NK T cells (n=2131) in light blue, macrophages (n=652) in olive green, NK cells (n=385) in orange, proerythroblast (n=464) in coral red, plasma cells (n=559) in light green and dendritic cells (n=249) in red). (B) Cell type percentages from male mice present in each age group of the dataset are shown as a line plot. (C) The number of DZ and DE miRNAs detected in each cell type separated by up and downregulated genes are shown (dark green and light green respectively). (D) Heatmap showing top ranked miRNA genes that were differentially detected (DZ) across main cell types in the spleen (Fisher’s exact test associated p value < 0.05, absolute log2 FC>0.5). Fold changes between old and young cells are shown in each cell type. (E) Dot plot heatmaps of myeloid-specific changes in miRNA gene levels (adjust p value <0.05). (F) UMAP plots of macrophage and dendritic cell gene markers (on the left) and cell type-specific miRNAs (on the right).

Several previously aging-associated miRNAs are among the top ranked upregulated miRNA genes in old mice, including elevated expression of mmu-mir-101a, in agreement with its established role in aging brain tissue ^18^, and mmu-mir-146a and mmu-mir-147 that regulate NF-κB and Toll-like receptor (TLR) mediated inflammatory responses and induce pre-mature senescence ^19–21^. In our analysis, mmu-mir-146a and mmu-mir-147 were concordantly upregulated in the immune cell types analysed and mmu-mir-101a expression increased most in lymphoid cells (T-cells, B-cells, and NK T-cells) of old mice (Fig. 2D, top). Interestingly, female samples comprising young (3 months) and adult mice (18 and 21 months) only showed common upregulation of mmu-mir-147 and decrease of mmu-mir-146a and mmu-mir-101a (Fig. S2C).

Further expression changes seen in aged immune cells that may aggravate aging phenotypes include downregulation of mmu-mir-706 (Fig. 2D, bottom) with recognized function as an oxidative stress regulator ^22^, upregulation of mmu-mir-30a∼30c-2 and mmu-mir-30b∼30d (upregulated in 4/5 cell types) representing members of miR-30 family microRNAs that promote senescence (inhibition of DNA synthesis by targeting B-myb ^23^), the concordant increase in mmu-mir-705 (regulation of aging-related cell fate bias ^24^) and mmu-mir-691 that may promote chronic inflammation (ulcerative colitis ^25^).

Our analysis also identified highly cell type-specific changes, exemplified by a decrease in mmu-mir-455 transcription in myeloid cells in line with aging-protective function in osteoarthritis ^26^ (Fig. 2C, E and Fig. S2D). Similarly, the fraction of cells expressing mmu-mir-511 involved in the regulation of TLR-signalling decreased. In contrast, we observed upregulation of mmu-mir-138-1 and mir-676 transcription towards aging in myeloid cells. Previously similar pattern towards aging has been reported in keratinocytes were miR-138 promotes cellular senescence via targeting *Sirt1* ^27^. These myeloid-lineage specific changes matched specific sub-populations of cells, corresponding to dendritic cells and macrophages (refer to Fig. 2F showing the respective marker genes: *H2-DMb1* for antigen presenting dendritic cells and *Cd163* for macrophages).

Taken together, our analysis revealed aging-related transcriptional changes of miRNA genes involved in regulatory networks governing senescence, oxidative stress and inflammatory responses, distinguishing several miRNAs impacted in multiple immune cell types and providing the resolution to detect highly cell type-specific expression.

### Identification of miRNA gene markers for myeloid subpopulations in fat tissue

An unhealthy diet is a risk factor for disease development that can result in elevated white adipose tissue (WAT) inflammation through altered cytokine and chemokine secretion in which specific immune cells are key players ^28^. To model this process and analyse changes in miRNA gene expression, we collected scRNA-seq profiles from conditions representing progressive atherosclerosis development in LDLR^-/-^ApoB^100/100^ mice fed with a chow or high fat diet (n=5 per group). The experimental setup led to atherosclerotic plaque formation phenotype resembling early disease state (ED, shorter fat diet) and late disease (LD, longer fat diet resulting in advanced vascular lesions) (Fig. 3A), confirmed by examining the vessel wall cell phenotypes at lesions ^29^. In addition, we included lipopolysaccharide (LPS) as an extra inflammatory stimulus introduced during the fat diet (two weeks prior to tissue collection in ED condition) to achieve an inflammatory challenged state (IC, n=5) (Fig. 3A). In response to IC and also at LD, the proportions of immune cells (T cells, B cells, and myeloid cells shown in Fig. 3B) were modulated, with concomitant decrease in relative proportion of non-immune tissue-resident stromal cells. Among the immune cell types, the myeloid cell fraction increased the most between different conditions.

**Figure 3.**
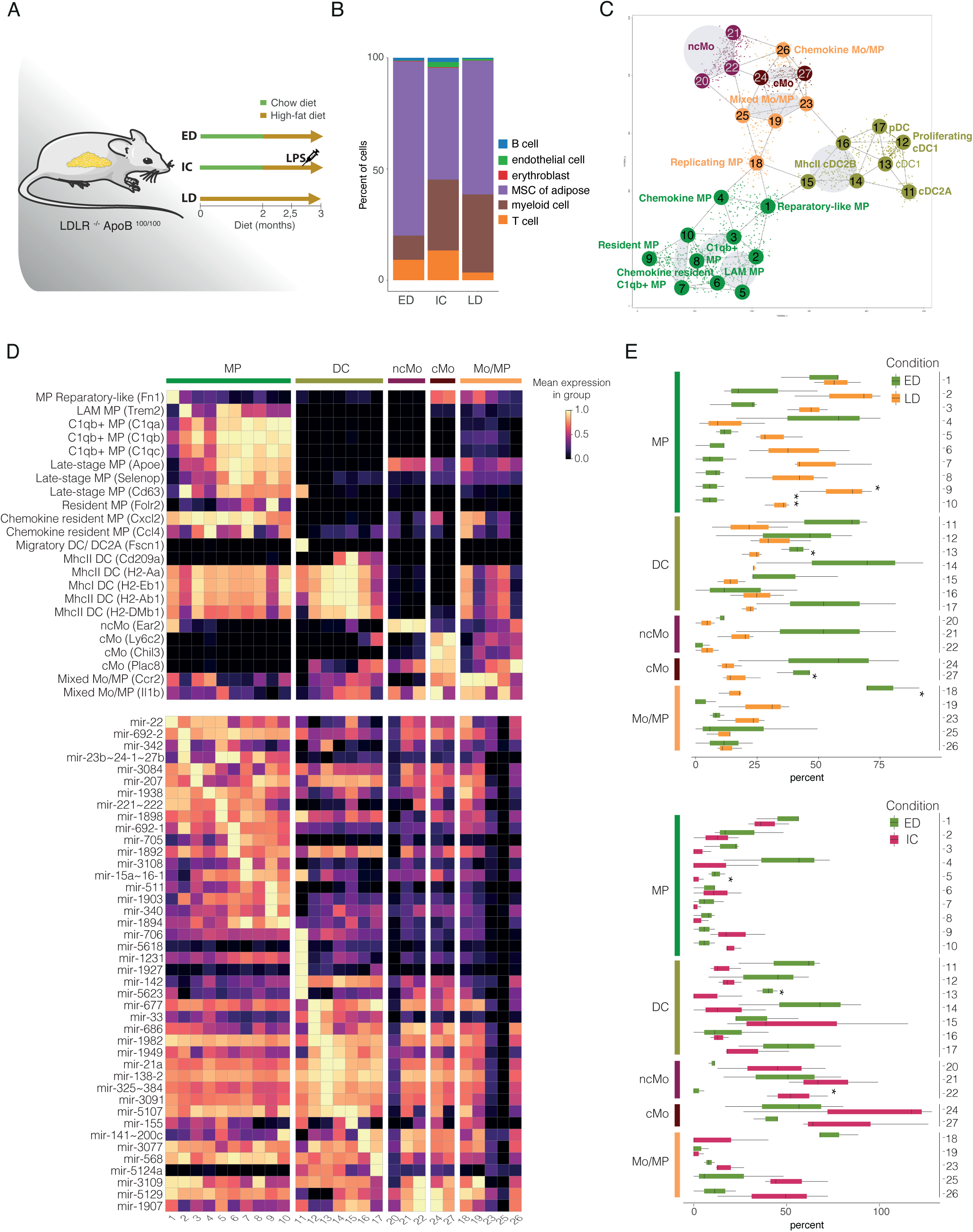
Characterization of myeloid cell subpopulations and their miRNA gene profiles during atherosclerosis disease progression. (A) Scheme of the study of atherosclerotic plaque formation in mice. Mice were fed either a chow diet (in green) or fat diet (in gold). Samples were denoted as early-disease (ED), inflammatory challenge (IC) and late-disease (LD) for LDL deficient mice. (B) Proportion of cell types across the sample types. (C) Cells grouped in meta nodes based on their reproducible phenotypes. (D) Heatmap showing expression of selected node marker genes (above) and marker miRNA genes (below). Expression is row-scaled. (E) Changes in cell proportions between ED and LD conditions (above) and between ED and IC conditions (below).

To characterize further the myeloid compartment, we defined reproducible cell phenotype states by computing a cell similarity graph from the scRNA-seq profiles using the MetaCell pipeline ^30^. To annotate the 27 nodes found that each represent a unique transcriptome state (Fig. 3C), we obtained marker genes and visualized their expression alongside the expression of literature-based markers (Fig. 3D, Fig. S3A, Table S3A).

Macrophage (MP) markers were highly expressed in Nodes 1-10. Specifically, node 1 had elevated expression of genes such as *Retnla* and *Fn1* (termed the reparatory-like MPs). Node 2 showed elevated expression of *Trem2* that is a well-known marker of lipid-associated macrophages (termed the LAMs) ^31^. Nodes 3 and 5-10 showed a higher expression of genes encoding for innate immune protein *C1q* expression that in macrophages was previously suggested to alleviate inflammation present during atherosclerosis disease progression ^32^. Accordingly, cells matched to nodes 5-10 predominantly represented LD condition and showed a relatively higher expression of late-stage MP markers (*Apoe*, *Selenop*, and *Cd63* ^33^). In addition to node 2, nodes 5 and 6 had elevated expression of *Trem2*, indicating a LAM-like transcriptional signature. Node 9 showed a high expression of *Cd163*, *Lyve1*, and *Folr2* (termed as the tissue-resident MPs ^33^). Interestingly, nodes 4 and 7 were enriched in genes associated with chemokine-signaling such as *Cxcl2* and *Ccl4*, suggesting that node 4 is composed of chemokine MPs and node 7 compose of *C1q*+ MPs with also chemokine secretion. Additionally, node 7 showed an intermediate expression of *Cd163* and *Folr2* further implying a tissue-resident-like MP phenotype.

Node 11 showed a distinct transcriptional state with a high expression of *Ccr7*, *Ccl22*, and *Fscn1* (Fig. 3D, Fig. S3A). Cell populations enriched with these markers have previously been termed the migratory dendritic cells (DC) ^33^ and the classical DC2A ^34^. *Xcr1* and *Clec9a* were high expressed only in nodes 12 and 13 suggesting cDC1 phenotype ^33,35^. Additionally, node 12 was enriched in genes encoding the members of Cdc45/Mcm2-7/GINS (CMG) complex (*Mcm5* and *Mcm6*); thus, this node was termed as proliferating cDC1. Nodes 14, 15, and 16 had elevated expression of *Cd209a* together with major histocompatibility complex (MHC) II class genes (*H2-Eb1*, *H2-Ab1*, *H2-Aa*, *H2-DMb1*) and therefore defined as MHCII DC ^33^. Plasmacytoid DC markers (*Siglech*, *Ccr9*, *Cox6a2*, *Atp1b1*, *Ly6d* ^33^) were exclusively expressed in node 17.

Node 18 showed a mixture of different signatures: monocyte-derived MP (*Ccr2*), interferon (INF) (*Isg15*), chemokine (*Cxcl10*), and active DNA replication (*Top2a*) suggesting that they are actively replicating MPs undergoing transition potentially towards INF or chemokine MPs (Fig. 3D). Markers of mixed Mo/MP (*Ccr2* together with *Fcgr1*, and *Itgam* ^33^) were present in node 19. Since nodes 20-22 showed upregulation of *Ace* and *Ear2* together with downregulation of *Ccr2* and *Ly6c2*, they were defined as non-classical monocytes (ncMo ^33^) (Fig. S3A). In contrast, nodes 24 and 27 showed upregulation of *Ccr2* and *Ly6c2* together with high expression of *Chil3* and *Plac8*, and these nodes were thus defined as classical monocytes (cMo ^33^). Nodes 23 and 25 showed mixed patterns of Mo and MP such as intermediate expression of *Ccr2* (Mo-derived MP), *Isg15*, *Isg20* (INF), *Fcgr1*, and *Itgam* (Early MP) (Fig. 3D). Node 26 was enriched in genes associated with INF signatures such as *Isg15* and *Isg20* and was therefore defined as Mixed Mo (Fig. S3A).

Comparison of relative cell proportions revealed that macrophage nodes 1-3, 5-10, 19, and 23 had high representation of cells in LD condition whereas ncMo nodes 20-22 and cMo/Mo-MP nodes 24-27 were predominant in the IC condition (Fig. 3E, Fig. S3B). We next examined miRNA gene expression specific to the metacell subpopulations. Highly node-specific expression of several immunomodulatory miRNA genes distinguished the macrophage subtypes, in comparison to more subtle subtype-level differences between DC and monocyte (Mo) cell subtypes (Fig. 3E). The macrophage miRNA markers included highest mmu-mir-22 levels (regulation of proinflammatory cytokine expression ^36^) within reparatory MP, LAM-specific expression of mmu-mir-23b∼24-1∼27b (regulation of proinflammatory cytokine expression, downregulated miRNA in patients with autoimmune diseases ^37^), high mmu-mir-221∼222 in *Cq1*+ MP and mmu-mir-15a∼16-1 (regulating phagocytosis ^38^) in MP node 7. The most distinct miRNA gene expression among DC was found in cDC2 (e.g. mmu-mir-706 with possible unconventional nuclear function ^39^) and in n15 cells corresponding to MHCII DC phenotype, that express mmu-mir-155, known to function as a “master regulator miRNA” in DCs and MPs ^40^. In DCs mmu-mir-142a has key role in regulation of proinflammatory cytokines in DCs ^41^. In agreement, our analysis identified it as a broadly expressed DC marker miRNA.

### Disease progression alters miRNA gene expression in macrophage subpopulations

Next, we compared the miRNA gene expression distributions in ED and LD conditions in myeloid cells. Across all cells, our analysis identified 21 upregulated and 6 downregulated miRNA genes (Fig. 4A, Fig. S4A, Table S4). Among immunomodulatory miRNAs with potential to aggravate tissue inflammation, we noted increased expression of mmu-mir-511 ^42^ driven by nodes 2-8 (Fig. 4A). An opposite change was observed for mmu-mir-101b expression (negative regulator of pro-inflammatory response ^43^, with strongest repression in nodes 23 and 25 (nodes panel, Fig. 4A). In comparison, immunosuppressive mmu-mir-23b∼24-1∼27b expression increased in Trem2 and Trem2-like MPs (nodes 2 and 5) (nodes panel, Fig. 4A) and miRNA genes encoding classical nuclear factor kappa B (NF-κB)-modulating miRNAs mmu-mir-146a and mmu-mir-21 ^42^ increased in expression in several MP nodes, mmu-mir-146a most strongly in node 7. Taken together, our results demonstrate that disease development strongly modulates miRNA gene expression and that these changes include both pro- and anti-inflammatory regulatory pathways with distinct expression across macrophage subtypes.

**Figure 4.**
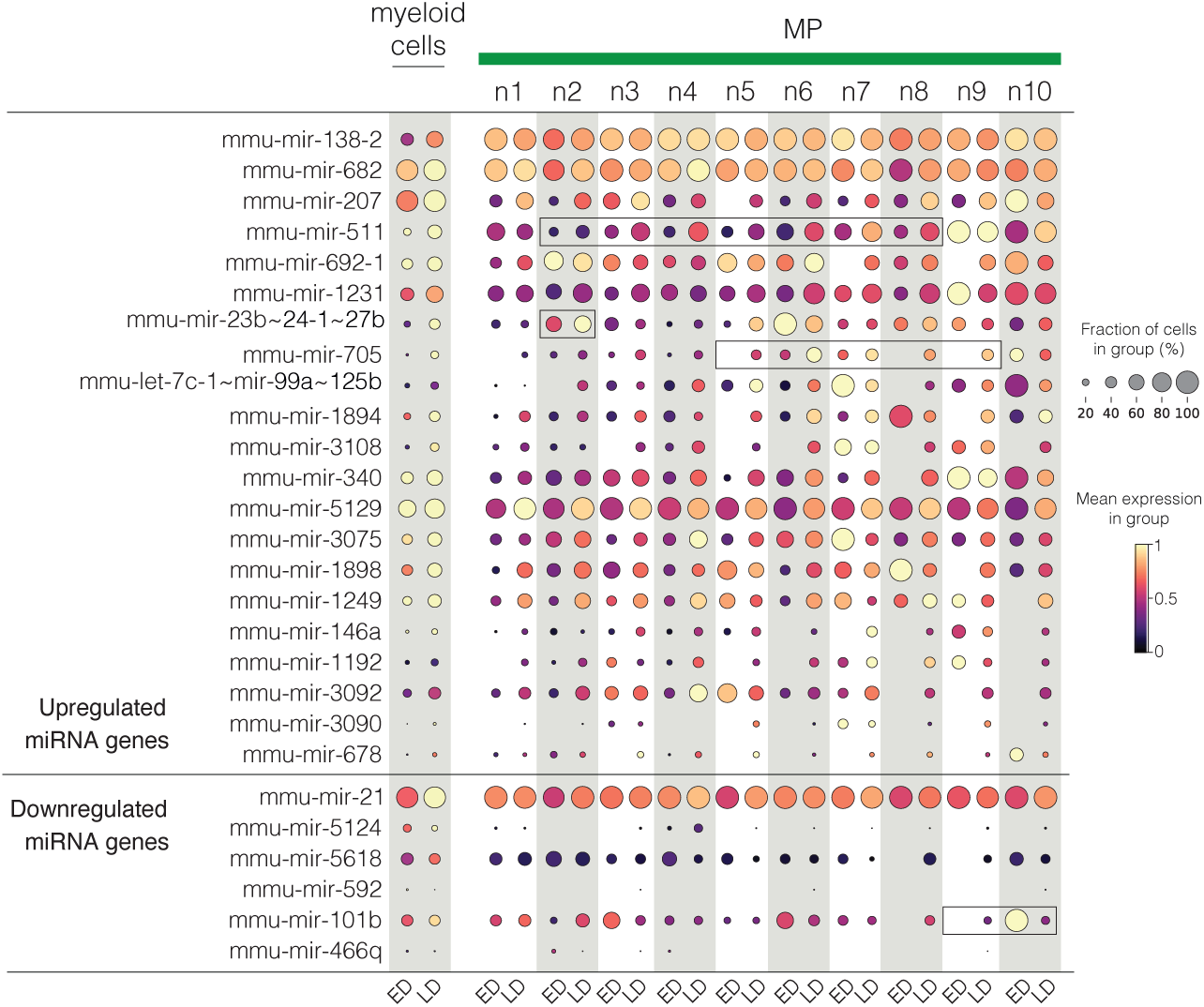
Disease progression alters miRNA gene expression in macrophage subpopulations. miRNA genes with altered expression during disease progression are shown as dot plot heatmap and stratified by metanode. Brighter color tone and larger dot size denote higher expression.

### Tissue-infiltration of monocytes and transition into mature phenotypes strongly modulates miRNA gene expression

The prominent increase in myeloid cells in adipose tissue at late disease, or at early disease upon LPS-stimulus, prompted us next to examine changes in gene regulation during myeloid maturation into tissue-resident cells. We hypothesized that the cytokine environment within tissue could trigger changes in gene regulation and thereby miRNA expression. Thus, we collected additional scRNA-seq profiles from blood monocytes and integrated these with the myeloid cell profiles from WAT. The monocytes in blood are typically short-lived, representing a reference naïve state for the comparison. The cells obtained from blood and WAT clustered primarily based on their tissue-of-origin (Fig. 5A, UMAP), however with similar sub-populations (ncMO; cMO; DC) from both tissues placed adjacent to each other, as defined using marker genes (Fig. S5A, see also Fig. 3 and Fig. S3). We focused on the main monocyte and DC subtypes and performed statistical comparison of their tissue vs blood expression profiles (Fig. 5B, Table S5A-B, refer also to Fig. S5B for DC comparison). In total, our analysis detected significant changes in 45 and 95 miRNA genes in ncMo and cMo, respectively. Among the top upregulated miRNA genes, mmu-mir-1938 and mmu-mir-22 are highly upregulated in both monocyte types, while the most significant changes in cMo (more pro-inflammatory monocyte type) include also upregulation of mmu-mir-221∼222, mmu-mir-511, and mmu-mir-155. The comparison of top miRNAs and classical LPS-responsive genes (*Dusp1*, *Il1b*, *Ccl5*) in blood and WAT (Fig. 5C) highlights increased expression of these pro-inflammatory genes (dot size and darker red color tone) upon tissue infiltration that is further elevated in IC condition, most prominently in ncMo.

**Figure 5.**
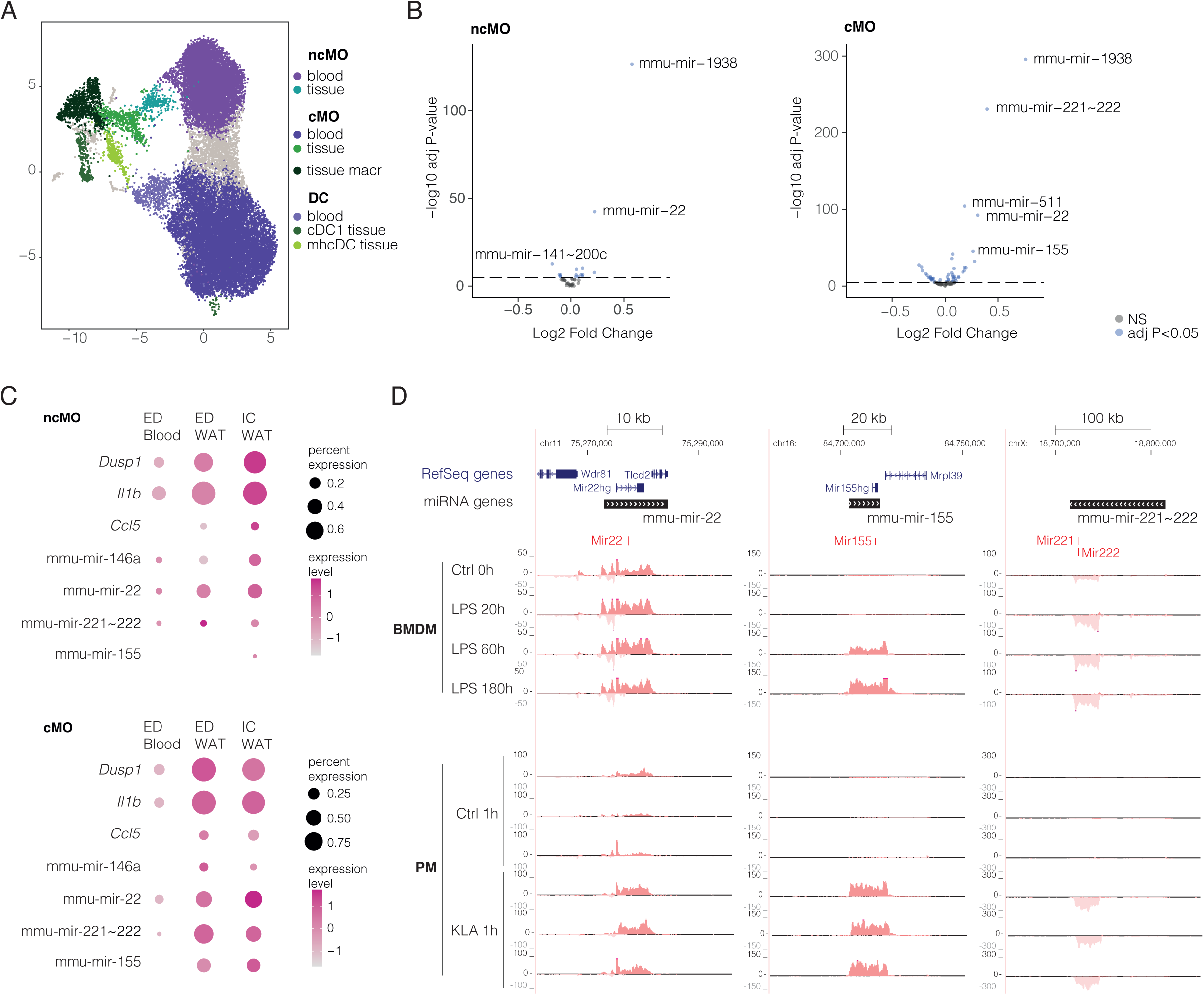
Gene expression changes in monocytes infiltrating adipose tissue relative to naïve blood precursors. (A) UMAP of blood and tissue (WAT) myeloid cell populations. (B) Volcano plots representing significant miRNA gene loci from tissue vs blood comparison of monocyte subtypes. Top miRNA loci are indicated on the plot. FC correspond to differences in detection rate. (C) Dot plot heatmap comparing known LPS-responsive genes to the profile of top miRNA loci in tissue and blood. GRO seq signal tracks at mmu-mir-22, mmu-mir-155 and mmu-mir-221∼222 loci. The + strand signal is shown above the vertical axis (dark red tone) and -strand signal below (light red tone). NcMo: non-classical monocyte, cMo: classical monocyte, DC: dendritic cell, ED: early disease, IC: inflammatory challenge, BMDM: bone-marrow derived macrophage, PM: peripheral macrophage, LPS: lipopolysaccharide, KLA: Kdo2-lipid A).

To validate that the changes detected from scRNA-seq profiles represent regulation of the transcriptional activity at pri-miRNA loci, we used GRO-seq datasets (see Methods) collected from two different experimental setups: *ex vivo* LPS stimulation of bone-marrow derived CD14+ macrophages (referred to as BMDM) and LPS stimulation of peritoneal MPs (referred to as PM, resembling tissue-resident MP). Three intergenic miRNA gene loci highlighted in the scRNA-seq analysis (Fig. 5A and B) are shown in Fig. 5D. The elevated GRO-seq signal levels within the gene regions confirm that upregulation of mmu-mir-22 and mmu-mir-221∼222 transcription occurs rapidly in both BMDM and PM cultures and remains high 180 h after LPS stimulus (see Table S5C-D for differential expression statistics and summary). In comparison, mmu-mir-155 is significantly upregulated with delay at 60 h in BMDM, representing a more immature cell model. In peritoneal cells its gene regulatory dynamics are more rapid (upregulation at 1 h) and comparable to the two other miRNA loci, overall in agreement with their increased expression in the tissue microenvironment and following LPS-stimulation in the *in vivo* scRNA-seq profiles.

## DISCUSSION

MiRNAs are key modulators in maintaining tissue homeostasis by post-transcriptionally regulating gene expression. Thus, an imbalance in their expression has been associated with disease progression and aging-related cellular processes including senescence ^44,45^. Here, we aimed to characterize miRNA gene transcription in single-cell datasets to study cell-type specific miRNA expression. We provided primary transcript annotations for mouse and human genomes that can be added to scRNA-seq quantification pipelines and demonstrated feasibility to capture miRNA gene expression from both droplet- and plate-based platforms. Based on this approach, we quantified and analysed miRNA gene expression during aging in splenic immune cells and delineated changes in myeloid cell populations upon atherosclerosis disease progression in blood and adipose tissue. The identified changes in miRNA gene expression at cellular resolution include well-established miRNAs that control inflammation, senescence, and metabolic responses in immune cells, supported by previous functional studies and analysis of nascent transcription in *ex vivo* cell culture models. We provide the miRNA gene profiles across the aging and atherosclerosis time series as an openly available resource to facilitate further characterization of miRNA gene expression changes related to tissue homeostasis.

In this study, we extended reference transcript annotation with miRNA gene coordinates and evaluated how two commonly used scRNA-seq platforms serve in miRNA gene detection and cell subpopulation identification. The larger number of cells profiled using the 10x Genomics allows a better estimation of the heterogeneity of a cell population ^7,46^. In line with previous results, the TMS 10x Genomics profiles captured a higher proportion of non-coding transcripts compared to Smart-seq2 data ^7^. Plate-based studies, on the other hand, often aim at higher sequencing depth per cell and have full gene body signal coverage. This could benefit the characterization of differences in alternative transcripts within miRNA gene loci. However, if strand specificity is lacking, this analysis has similar limitations as chromatin immunoprecipitation (ChIP)-seq based analysis of transcriptional activity, described in ^10^. In both platforms, the detection of miRNA gene transcription suffers from limited number of intronic reads captured. In future, as more nascent transcriptome single cell profiles become available, better capture of miRNA gene expression levels could be achieved. Among existing data, scRNA-seq profiles generated from nuclei already provide higher capture of intronic reads ^47^ that will improve miRNA gene detection ^48^. As the signal within pri-miRNA gene loci in standard scRNA-seq libraries can arise from random priming at unspliced introns or polyA-based capture of incompletely processed primary transcripts, we chose to base the miRNA gene annotation on integrated GRO-, CAGE- and ChIP-seq ^9^ and Drosha knockout (KO) profiles ^49^, extending here to cover both human and mouse genomes. Alternative transcript assembly-based approach ^50^ can leverage the data only partially, and as key limitation its feasibility is impacted by insufficient cell numbers per cell type. Furthermore, quantification of miRNA genes based on the pre-defined coordinates we provide here extends to other data types, as it can be readily introduced into single-cell sequencing assay for transposase-accessible chromatin (scATAC-seq) and scMultiome data analysis pipelines to analyse chromatin accessibility at miRNA loci, enabling the identification of their corresponding regulatory elements residing in open chromatin. Previous studies extending to chromatin signatures have provided key new insight on cell type-specific gene regulatory network activity focusing on transcription factors ^51^. The joint analysis of miRNA genes will enable a more comprehensive analysis of gene regulatory networks that govern tissue homeostasis at transcriptional and post-transcriptional levels.

Aging is a stepwise process characterized by changes in tissue homeostasis and cellular heterogeneity. For example, changes in adipose depots distribution along the body dramatically affect tissue growth, plasticity and function leading to metabolic dysfunction and low-grade inflammation ^52^. Cross-talk between adipose tissue and immune cells is crucial for the maintenance of normal healthy adipose tissue function and systemic metabolism ^53^. Consequently, better understanding of the miRNA post-transcriptional regulatory networks that fine tune cytokine expression and response dynamics upon inflammatory challenge can provide new approaches to predict and prevent progression of age-related functional changes ^54^. Mature miRNAs are highly stable and therefore ideal biomarker candidates for monitoring disease progression. Moreover, new therapeutic approaches based on miRNA-delivery into tissues are in development, with promise to reduce the burden of aging and immune dysfunction-related disease including type 2 diabetes, atherosclerosis, dyslipidaemia, thermal dysregulation and skin ulcers among others ^55^. In this study, we performed comparisons of miRNA gene expression in immune cells from different tissues, including bone marrow-derived blood cells and more mature splenic and tissue-resident populations. Our analysis identified altered expression of several miRNA genes (mmu-mir-101, mmu-mir-30, mmu-mir-709) in spleen that have established function in regulation of immunosenescence and apoptosis that throughout the body presents with alterations of immune cell homeostasis and an overall decline in immune efficacy ^56^. The TMS atlas affords opportunity to extend these comparisons to additional tissues. We limited the statistical comparisons to male mice, which is different from earlier comparisons that included both sexes ^13^. Our more conservative choice relates to the lack of representation of both sexes in certain age groups, with more timepoints available in male mice with sufficient time difference ^13^. For example, in spleen tissue our analysis of male data included young (1 and 3 months) versus old (21 and 30 months) comparison, while the female data corresponded to adult (18 and 21 months) versus young (3 months) comparison. Concordantly, the profiles from our atherosclerotic mouse model represent male mice, enabling comparison to TMS data without confounding sex effect. The different aging profile of mmu-mir-146a in females suggests that miRNA gene regulation is impacted by sex and is in agreement with a recent human study reporting that the miR-146a age-related trajectory was confirmed only in men ^57^. Therefore, in future studies it would be important to collect data representing more comprehensively both male and female aging. More broadly, sex-biases in immune responses are well-established and known to strongly influence disease prevalence ^58,59^. As the technologies mature, integrating cell-specific mature miRNA profiles will expand the analysis of miRNA genes from the TMS and similar aging atlases, leading to better understand the functional impact of miRNA dysregulation that can account for concomitant changes in RNA processing that occur during aging ^60,61^, including decreased levels of the miRNA processing enzyme Dicer both in mouse and human ^62^. Furthermore, analysis of immune cell types at finer resolution should be pursued, as was carried out here for myeloid cell subtypes, guided by updates to atlas cell type annotations and improved understanding of the functional differences between the new cell subtypes identified from single cell studies.

Traditionally, macrophages were categorized as pro-inflammatory (M1) and anti-inflammatory (M2) macrophages that show distinct miRNA profiles, for example miR-511 expression is increased in M2 and decreased in M1 macrophages both *in vitro* and *in vivo* ^63^. However, several studies have suggested this dichotomy to be obsolete and that M1 and M2 stages rather represent the extremes of a spectrum in a multidimensional space ^64,65^. Recent single cell studies have characterized macrophage subpopulations within the fat tissue, revealing subtypes involved in obesity disease progression, including the description of Trem2+ lipid-associated macrophages ^34^. Similarly, single cell profiling of alveolar macrophages revealed novel macrophage subdivisions based on proliferation capacity and inflammatory programming in the context of lung inflammation ^66^. Here, we conducted analysis guided by unbiased clustering of cells into nodes showing distinct transcriptional phenotypes. WAT myeloid subtypes differed in miRNA gene expression, with highest basal miR-511 levels in tissue-resident macrophages (nodes 9 and 10) and increased expression found across multiple subtypes upon disease progression. The highest expressed miRNA genes in Trem2+ macrophages were mmu-mir-221∼222 and mmu-mir-23b∼24-1∼27b loci. Previous studies have demonstrated a functional role for miR-221/222 in inhibiting adipogenesis and preventing diet-induced obesity. At systemic level, *Mir221/222AdipoKO* mice used in these studies did not show significant improvement of insulin sensitivity. Instead, lower expression of these genes may promote apoptosis upon hyperglycaemia ^67^. Furthermore, mmu-mir-23b∼24-1∼27b locus from which miR-23b, miR-27b, miR-24-1 originate from encodes miRNAs that each have a central role in regulation of lipid metabolism ^68–70^. Thus, changes in their expression could strongly impact the lipid-associated functions of the Trem2+ cell phenotype. Moreover, a recent study that generated a specific mouse KO of this miRNA locus showed impaired glucose tolerance ^71^, emphasizing the critical role of miRNAs in maintenance of homeostasis. Here we observed that upon inflammatory challenge (LPS stimulation) mmu-mir-221∼222 expression increased, while mmu-mir-23b∼24-1∼27b and *Trem2* levels decreased in the Trem2+ macrophage. Thus, future studies examining the connection between Trem2-specific cell-intrinsic regulatory networks and systemic glucose and lipid metabolism balance are warranted.

Infiltration of immune cells, especially those with pro-inflammatory function, into tissues is a known hallmark of aging ^52^. We found that several key miRNA genes were regulated upon monocyte recruitment into tissues. Among them, miR-155 induces pro-inflammatory activation of monocytes and through increased expression of human leukocyte antigen (HLA)-DR in the myeloid cells, it also modulates activation of T cells and can thereby aggravate tissue inflammation, promoting disease progression in inflammatory-disease such as arthritis ^72^. However, the induction of mmu-mir-155 coincided with elevated expression of miRNA genes typical of M2-like cells, such as miR-146a and miR-511 ^73^. In addition, our analysis identified several less well-characterized miRNAs, including mmu-mir-1938, that merit future functional characterization. The top target prediction for miR-1839 is *Laptm5* (TargetScanMouse 7.1), encoding a lysosomal protein that modulates pro-inflammatory signalling in macrophages ^74^. Elucidating the mRNA targets using immunoprecipitation of argonaute family members followed by RNA sequencing or reporter-based screens ^75^ can provide a more comprehensive understanding of how the miRNAs that are transcriptionally regulated upon tissue entry collectively impact tissue cytokine environment, immune-cell interactions, and tissue homeostasis.

In summary, our work provides an unbiased and genome-wide evaluation of miRNA loci at cellular resolution during aging and upon disease development and new tools for single-cell genomics research to define miRNA regulatory networks with coordinated and cell type-specific activities across tissues.

## MATERIAL AND METHODS

### Annotation of miRNA gene coordinates in mouse genome

To distinguish primary transcripts corresponding to miRNA genes in mice, we followed the strategy we introduced in ^9^. To define mouse pri-miRNA genes, GRO-seq and CAGE-seq data aligned to the mm9 genome were used to define genomic intervals that correspond to active primary transcription, separated by active TSS. Due to fewer mouse GRO-seq samples available, the known RefSeq and UCSC known gene (2018) transcript data ^76,77^ was included as external data to complement the *de novo* transcript discovery. TSS coordinates were refined based on CAGE-seq, while the extension of transcript ends was defined based on signal change point analysis from GRO-seq, or if the available annotated gene region matched the candidate transcript the longer transcript region between the annotated and discovered transcribed region was kept. Finally, the pre-miRNA annotations from GenCode v19 and miRbase v.20 ^60^ were used to annotate the subset of primary transcripts that overlapped miRNA coordinates. The coordinates were then converted to mm10 using UCSC liftOver tool ^78^ to be compatible with the most recent genome version.

### Building a custom transcript annotation for scRNA-seq gene quantification workflows in mm10 and hg19 genomes

Typical scRNA-seq quantification workflows, including the 10x Genomics Cell Ranger pipeline ^79^, allow users to build their own custom transcript annotation based on existing reference annotation data that describes gene, transcript and exon information. To develop a custom reference suitable to quantify the transcriptional activity of miRNA genes, we defined miRNA genes as the merged region starting from the most distal TSS mapped to the miRNA and extending until the longest transcript end, using the GRO- and CAGE-seq-based annotation for previously defined human coordinates^9^ and mouse coordinates described above. This region was included in the GTF file as a single exon (pri-miRNA) transcript. However, also alternative transcript and exon structures have been experimentally defined for miRNA genes based on knockout of the key processing enzyme Drosha ^49^. Therefore, the annotation was extended by adding these candidate alternative transcript structures and at annotated coding gene regions exons were included from GenCode (2018 and 2013 for mouse and human genomes respectively), motivated by manual examination of splicing patterns captured in 10x Genomics and Smart-seq2 scRNA-seq data. Cell Ranger requires that GTF files are preformatted using ‘cellranger mkgtf’ command and that a FASTA file (reference genome) containing the nucleotide sequences of the selected transcripts is provided. The generated GTF with miRNA genes and the FASTA file for mouse mm10 or human hg19 genome were used as input for the ‘cellranger mkref’ command. The additional quantification for the miRNA genes was combined in downstream analyses with the default GenCode-based count matrix.

### TMS and Tabula Sapiens Data

The TMS and Tabula Sapiens consortia generated single-cell libraries that were produced either using single-cell suspensions combined with droplet detection (10x Genomics) or by fluorescence-activated cell sorting (FACS) sorting individual cells combined with Smart-seq2 technology hereafter referred as “Plate-seq” as denoted in the original TMS publications ^11–13,15^. In this work, we used data from spleen. However quantified data is shared for Liver, Heart and Aorta, Fat and Bone Marrow tissues at 1 month, 3 months, 18 months, 21 months, 24 months, and 30 months of age for the tissues that were available. Selected files were downloaded from the amazon cloud as described in the GitHub repository available for this work (https://github.com/anahSG/scMIR/). A list of samples used can be found in Table S1D-E.

10x Genomics datasets were downloaded in .fastq format and processed with the default Cell Ranger pipeline (*v.3.0.2).* The detection of cell-containing droplets in 10x Genomics data was performed using the default count matrix, and the miRNA gene counts were added based on matching cell barcodes and quantified with the option –include-introns (v.6.1.1). Smart-seq2 samples denoted as ‘Plate-seq’ were downloaded in bam format and quantified with FeatureCounts (Subread package v.2.0.1, ^80^) using the custom GTF reference genome with default options (in this case, the libraries are strandless).

Droplets assigned by the Cell Ranger pipeline as cell-containing droplets (filtered matrix) were further quality controlled, filtered by QC metrics and processed using a standard SCANPY-based workflow described in ^17^ with minor modifications.

### Animal samples

LDLR ^-/-^ApoB^100/100^ transgenic mice have a phenotype characterized by high accumulation of fat in the tissues as they lack the ability to remove circulating lipid particles using the low-density lipoprotein receptor (LDLR) ^81,82^. This model is commonly used to follow atherosclerotic plaque formation in veins and arteries. To model disease progression, the transgenic mice were fed with a combination of chow diet and after, fat diet (HFD; 0.2% total cholesterol, Teklad TD.88137) for one month, capturing early-disease (ED) state. To study the impact of elevated pro-inflammatory signalling on immune cell tissue infiltration during disease progression, mice following ED diet were injected with LPS during the HFD phase two weeks prior to sacrifice. Late-disease (LD) state was achieved by feeding a fat diet for three months ^29^. C57BL/6J mice fed with a chow diet were as an additional control for this study (see annotation of cell types). We timed diet-starting age to equalize the age at sample collection between all groups (8 months old). Throughout the study, mice were maintained on a 12-h light-dark cycle and had access to food and water ad libitum.

All animal experiments were approved by the national Project Authorization Board (permission number ESAVI/4567/2018) and were carried out in compliance with the EU Directive 2010/EU/63 on the protection of animals used for scientific purposes.

### Tissue harvest, cell dissociation, and TotalSeq antibody staining

Mice were anesthetized with isoflurane and euthanized by cervical dislocation. Cardiac puncture was carried out with 10 mL of ice-cold PBS supplemented with 20 U/mL heparin and the mice were place into ice for dissection. ∼200 mg of epididymal WAT of each mouse (n=3 for each condition) was minced and added to 2.5 µl of Miltenyi Adipose Tissue Dissociation solution (supplemented with BSA and HEPES) and incubated for 45 min at 37 °C on an end-over-end rotator. During dissociation, the tissue was triturated 3 times (at 25 min, 35 min and 45 min) to break up cell aggregates. Tissue samples were passed through a 30 µm cell strainer and washed with 3 ml of RPMI. Cells were centrifuged at 400 g for 8 min at 4°C and the pellet was resuspended in 300 µl of FACS buffer (PBS with 1% BSA). Red blood cells were lysed using 1X RBC Lysis Buffer, Multi-species (eBioscience #00-4300-54) by mixing 300 µl of cell suspension with 2.7 ml of ice cold 1X RBC lysis buffer and incubating for 3 min on ice. 2 ml of FACS buffer was added to normalize the buffer and the cells were collected by centrifugation at 400 g for 8 min at 4°C. After RBS lysis, the cells collected from each mouse were stained with Total-Seq Mouse Hashtag antibody-DNA conjugates (BioLegend) containing a unique barcode sequence according to the manufacturers’ recommendations. The cell pellets were resuspended with TotalSeq Mouse Hashtag Ab-DNA conjugate in FACS buffer. The cell suspension was incubated in ice for 15-20 min to allow for hashtag Ab binding. Afterwards, cells were washed two times with FACS buffer to removed unbound hashtag Abs. Dead cells were removed with Miltenyi Dead Cell Removal kit (Miltenyi Biotec #130-090-101) as described before (Örd et al., 2023). The cell pellets were resuspended in PBS containing 0.04% BSA and counted by hemocytometry with trypan blue staining. Cell viability was between 74% and 85%. For each condition, approximately 18,000 cells (pooled from 3 mice) were loaded into the Chromium Controller microfluidics chip (10x Genomics).

Blood samples for monocyte isolation were processed by first performing erythrocyte lysis by mixing aliquots of 500 µl of EDTA blood with 4.5 ml of ice-cold 1x RBC lysis buffer and incubating on ice for 3 min. Subsequently, cells were centrifuged at 500 g for 5 min at 4°C, supernatants were discarded, and erythrocyte lysis was repeated. The cells were washed with 5 ml FACS buffer, followed by staining with TotalSeq Mouse Hashtag Ab-DNA conjugates in FACS buffer for 15-20 min on ice. Following staining, the cells were washed once with FACS buffer and once with MACS buffer (PBS with 0.5% BSA and 2mM EDTA). After staining, the CD115+ monocytes were enriched using Miltenyi Biotec MicroBead kit (# 130-096-354) as described by the manufacturer. The stained cells from individual mice (three per condition) were pooled in 90 µl MACS buffer, combined with 10 µl of the FcR Blocking Reagent (Miltenyi Biotec), and incubated for 10 min at 4°C. 10 µl of CD115-Biotin conjugates were added and the suspensions were mixed and incubated for 10 min at 4°C and the cells were pelleted by centrifugation at 300 g for 5-10 min at 4°C. The supernatants were discarded, and the cell pellets were resuspended in 80 µl of MACS buffer. 20 µl of Anti-Biotin MicroBeads were added to the solution, mixed and incubated for 15 min at 4 °C. The cells were washed with 1-2 ml of MACS buffer and centrifuged at 300 g for 5-10 min at 4°C. The cells were resuspended in 500 µl of buffer. MS columns were placed in the magnet and the samples were passed through 30 µm cell strainers before applying them to the MS columns. All the subsequent steps were performed at 4°C. The columns were washed three times with 500 µl of MACS buffer. After removing the column from the magnet, 1 ml of elution buffer was added, and the cells were flushed out by firmly pushing the plunger into the column.

The cells collected from both tissues were centrifuged at 300 g for 5-10 min at 4 °C, resuspended in PBS containing 0.04% BSA, and counted by hemocytometry with trypan blue staining. Cell viability was between 74% and 85%. For each condition, approximately 30,000 cells (pooled from 3 mice) were loaded into the Chromium Controller microfluidics chip (10x Genomics).

### Library preparation, sequencing, and alignment

ScRNA-seq libraries were generated with the Chromium Single Cell 3′ v.2 assay (10× Genomics). Libraries were sequenced using the NovaSeq 6000 platform (Illumina) to a depth of approximately 300 million reads per library with read lengths of 26 (read 1) + 8 (i7 index) + 0 (i5 index) + 91 (read 2). Raw reads were aligned to the mouse genome (mm10) using Cell Ranger (count pipeline) (v.3.0.2).

### scRNA-seq data integration and label transfer in the atherosclerotic mouse model

Expression data was loaded in the Seurat R package v.4.0.0 for integration prior to cell type prediction. Data was normalized using SCTransform to account for differences in sequencing depth. The reference dataset used in integration was formed by the unbiased integration of the control (C57BL/6J on chow diet) and late disease conditions for each tissue. The rest of the conditions were then integrated using the canonical correlation (CCA) algorithm (k.anchor=20, dims = 1:50). Cell type predictions at broad lineage level were obtained by performing tissue-wide label transfer. WAT labels were transferred from the TMS fat dataset. Blood tissue labels were transferred from the 10x Genomics Peripheral blood mononuclear cell (PBMC) human reference dataset ^79^. Gene symbols were translated to mouse with the biomaRt R package v.2.54.1 ^83,84^.

### Data de-multiplexing by hashtag signals

The atherosclerotic mouse model samples were hashtag-barcoded with individual barcodes added to distinguish between mice. Tissues including WAT and aorta yielded initially very low signal in cells detected by Cell Ranger, resulting in ∼80% negative cells with default demultiplexing settings (mostly in cells that were annotated as stromal cells). To overcome this limitation, we used the DSB R library v.1.0.3 ^85^ that was built to estimate the difference between the actual antibody signal in cell-containing droplets and the background signal in empty droplets in cellular indexing of transcriptomes and epitopes (CITE)-seq single-cell libraries. The DSB workflow uses the raw matrices containing all the droplets available and produces a matrix of scaled protein signals vs the background signal. We used the hashtag scaled matrix as input to perform the hashtag based demultiplexing. We also noticed that the differences in the distribution of hashtag signal intensity between cells of distinct lineage (e.g. myeloid vs. lymphoid) were causing errors in the annotation of doublets across the tissue (proportions of negative cells or doublets would not typically be expected to vary by cell type). To overcome this, we performed the hashtag demultiplexing separately by the broad cell lineage annotation obtained using label transfer. We continued the analysis with cells that were annotated as individual cells (singlets) and repeated the sample integration.

### scRNA-seq quality filtering and normalization

To check the quality of the libraries generated, we followed a basic QC and filtering workflow using the SCANPY v.1.8.2 package for each tissue. Transcripts were filtered to include those that were present in more than 3 cells. To assess the viability of the cells, we quantified mitochondrial and ribosomal genes. Cells were filtered out according to a maximum mitochondrial gene expression percentage, a maximum number of counts and a minimum number of expressed genes. Due to heterogeneity in the raw data, these parameters were set for each tissue and condition (refer to TableS3).

Expression data was normalized with size factor values derived from data normalization using the scran R package v.1.26.2, and then log-transformed using the function scanpy.pp.log1p. Highly variable genes were calculated with scanpy.pp.highly_variable_genes, selecting the top 4000 genes for principal component analysis and dimensional reduction. Louvain and Leiden clustering at different resultions were performed to define similar transcriptome states that can be used for assigning lineage and cell type annotations.

### Differential expression analysis using scDD

The log2 counts based on vst transformation available in Seurat R package v.4.0.1 were used to compare differences in gene expression distributions in a given cell type between different sample groups (e.g. young and old, or late vs. early disease) using the scDD v.1.14 ^86^ package following the approach described in ^87^. This statistical analysis allows the detection of expression changes based on the fraction of cells expressing a certain transcript (DZ category) and among expressing cells by comparing the expression level (DE, DP and DM categories) ^86^. Transcripts with adj. p-value < 0.05 (Benjamini-Hochberg FDR method) were considered as significant.

Combined p-values were calculated based on the Fisher’s exact test (sumlog function from metap R package v.1.8) across all cell types to identify miRNAs that were concordantly regulated during aging. From those, miRNAs with associated p-values < 0.05 that passed a log fold change cut-off in at least two cell types were considered as top-ranking candidates presented in figures.

### MetaCell analysis of tissue myeloid cell sub-populations

Myeloid cells extracted from white adipose tissue samples were used to compute a cell similarity graph to obtain homogenous group of cells denoted as ‘metacells’ with the MetaCell R package v.0.3.7 ^30^. Gene-level statistics were computed with ‘mcell_add_gene_stat’ and featured genes were selected based on their variance (>0.8, ‘mcell_gsetfilter_varmean’) and number of UMIs (>100 UMIs in the entire dataset and selected genes are required to be detected in at least 3 cells with > 2 UMIs, ‘mcell_gset_filter_cov’). A balanced cell graph or ‘balanced K-nn graph’ was computed as previously described (K=100, ‘mcell_add_cgraph_from_mat_bknn)’. Next, we performed resampling (n=500) and generated the co-clustering graph (‘mcell_coclust_from_graph_resamp’, min. node size=20, cell partitions= 5.000 covering 75% of the cells). Metanodes assignment-derived statistics from the co-clustering step are evaluated with ‘mcell_mc_from_coclust_balanced’ command with default settings. To check that the metacell nodes are homogeneous, cells that highly deviate from their metacell’s expression profile were plotted as outlier cells (‘mcell_plot_outlier_heatmap’, data not shown) and filtered afterwards ‘mcell_mc_split_filt’. Gene markers per node were extracted for further analysis – ‘mcell_gset_from_mc_markers’. Metacells were projected into a 2D graph for visualization (‘mcell_mc2d_force_knn’,’ mcell_mc2d_plot’).

### Differential expression analysis of bulk GRO-seq profiles from macrophage *ex vivo* cultures

Samples listed in Table S1A were used for differential expression testing as previously described in ^9^. Low expressed transcripts were filtered and differences in transcription level between sample groups analyzed with limma and edgeR packages. Genes were assigned significant based on adjusted p-value < 0.05 (Benjamini-Hochberg method).

## AVAILABILITY

Data analysis code and links to .h5ad files comprising quantified scRNA-seq data objects with miRNA genes are available under GitHub repository (https://github.com/anahSG/scMIR/).

## ACCESSION NUMBERS

NGS data has been submitted to NCBI GEO data repository with the accession codes GSE241552: obebmimixdoztsv, (scRNA-seq from atherosclerosis mouse model), GSE241567: armhsaqyfxgzlmr (scRNA-seq in ST2 cell line), GSE241550: ololkqcgrxkfrax (new GRO-seq data generated for primir transcript detection). Other accession codes for datasets used in analysis are listed in Table S1A.

## SUPPLEMENTARY DATA

## ACKNOWLEDGEMENT

The authors wish to acknowledge UEF Bioinformatics Center (Biocenter Finland) and CSC – IT Center for Science, Finland, for computational resources; Turku single-cell omics core facility services (Biocenter Finland), Turku Bioscience and EMBL GeneCore sequencing team for sequencing service provided; Turku Center for Disease Modeling University of Turku TCDM (European Infrastructure for Translational Medicine) for animal housing. We would like to acknowledge researchers who shared their data in NCBI-GEO or public repositories and made it available. We gratefully acknowledge group members (Petri Pölönen, Juha Mehtonen and Minna Voutilainen) for assistance in miRNA gene analysis and Juha Kekäläinen and Buddika Jayasingha for assistance in data sharing.

## AUTHOR CONTRIBUTIONS

**AH de Sande**: Conceptualization, Data curation, Analysis, Methodology, Visualization, Writing the manuscript. **T Turunen**: Analysis, Visualization, Writing the manuscript. **M Bouvy-Liivrand**: Conceptualization, Experimental design, Sample collection, Methodology, Analysis, Supervision. **T Örd**: Design of the animal study, Animal work, Sample collection, NGS library preparation. **C Tundidor-Centeno**: Data curation, Analysis, Visualization. **H Liljenbäck**: Animal work, Sample collection. **S Palani**: Design of the animal study, Animal work, Sample collection. **J Virta**: Animal work, Sample collection. **OP Smolander**: Supervision. **A Roivainen**: Design of animal study, funding acquisition, Supervision. **T Lönnberg**: Design of the animal study, Sample collection, NGS library preparation, Funding acquisition. **MU Kaikkonen**: Design of the animal study, Supervision, Funding acquisition. **M Heinäniemi**: Conceptualization, Design of the animal study, Analysis, Methodology, Funding acquisition, Project administration, Writing the manuscript.

## Supplementary information

**Table S1. miRNA gene coordinates and related NGS dataset accession codes**

(A) The datasets used for defining genomic intervals for miRNA gene custom quantification and (B-C) ‘miRNA gene coordinates’ in mm10 and hg19 genomes. Related to Fig. 1. (D-E) ‘Dataset accession codes’ for 10x and SMART-seq2 ‘scRNA data used in this study. NCBI GEO accession codes are listed.

**Table S2. Spleen scRNAseq summary of differential gene expression across immune cell types analyzed**

Differential distribution analysis summary from comparison of old vs young male mice. The statistics for differential detection rate (DZ) (A) and expression level (DE) (C) of miRNA genes are provided separately for each cell type. Includes also ranking of miRNA based on Fisher test combined p-values across cell types (B and D for DZ and DE categories). Related to Fig. 2.

**Table S3. Atherosclerosis mouse model scRNA-seq data pre-processing settings and marker genes of WAT myeloid sub-populations**

(A) Marker protein-coding and miRNA genes for metanodes defined from myeloid sub-populations in WAT are listed. Related to Fig. 3. (B) Cut-off parameters used in quality filtering of scRNA-seq data collected from LDLR^-/-^ApoB^100/100^ mice.

**Table S4. WAT scRNAseq summary of differential gene expression in myeloid cells at late disease**

Differential distribution analysis summary from comparison of late disease vs early disease in LDLR-/-ApoB^100/100^ mice. The statistics for differential detection rate (DZ) (A) and expression level (DE) (B) are provided for myeloid cell comparison of miRNA genes. Related to Fig. 4.

**Table S5. WAT and blood myeloid cell scRNAseq and GRO-seq summary of differential gene expression**

Differential distribution analysis summary from comparison of tissue vs blood monocyte and monocyte-derived cell populations in LDLR-/-ApoB^100/100^ mice. The statistics for differential detection rate (DZ)(A) and expression level (DE)(B) are provided for miRNA genes. GRO-seq: DE analysis of ex vivo cultured bone marrow-derived CD14+ or peritoneal macrophage stimulated with pro-inflammatory LPS or KLA treatments. Related to Fig. 5.

**Figure S1.**
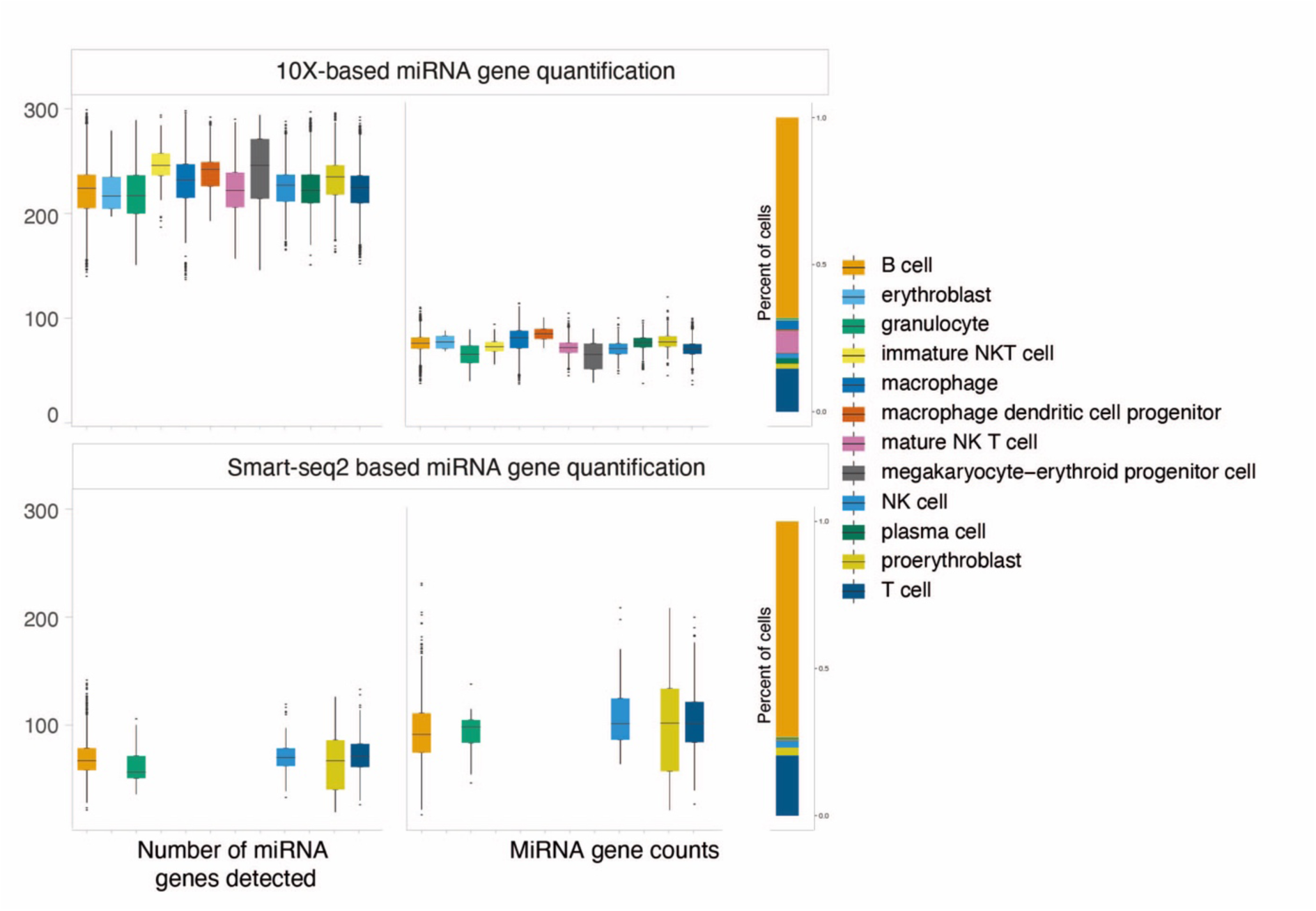
Comparison of miRNA gene expression measured from 10x Genomics and Smart-seq2 scRNA-seq technologies. MiRNA gene detection was evaluated in each splenic cell type by measuring the number of miRNA genes detected and their expression levels in 10x Genomics (upper panel) and Smart-seq2 technologies (lower panel).

**Figure S2.**
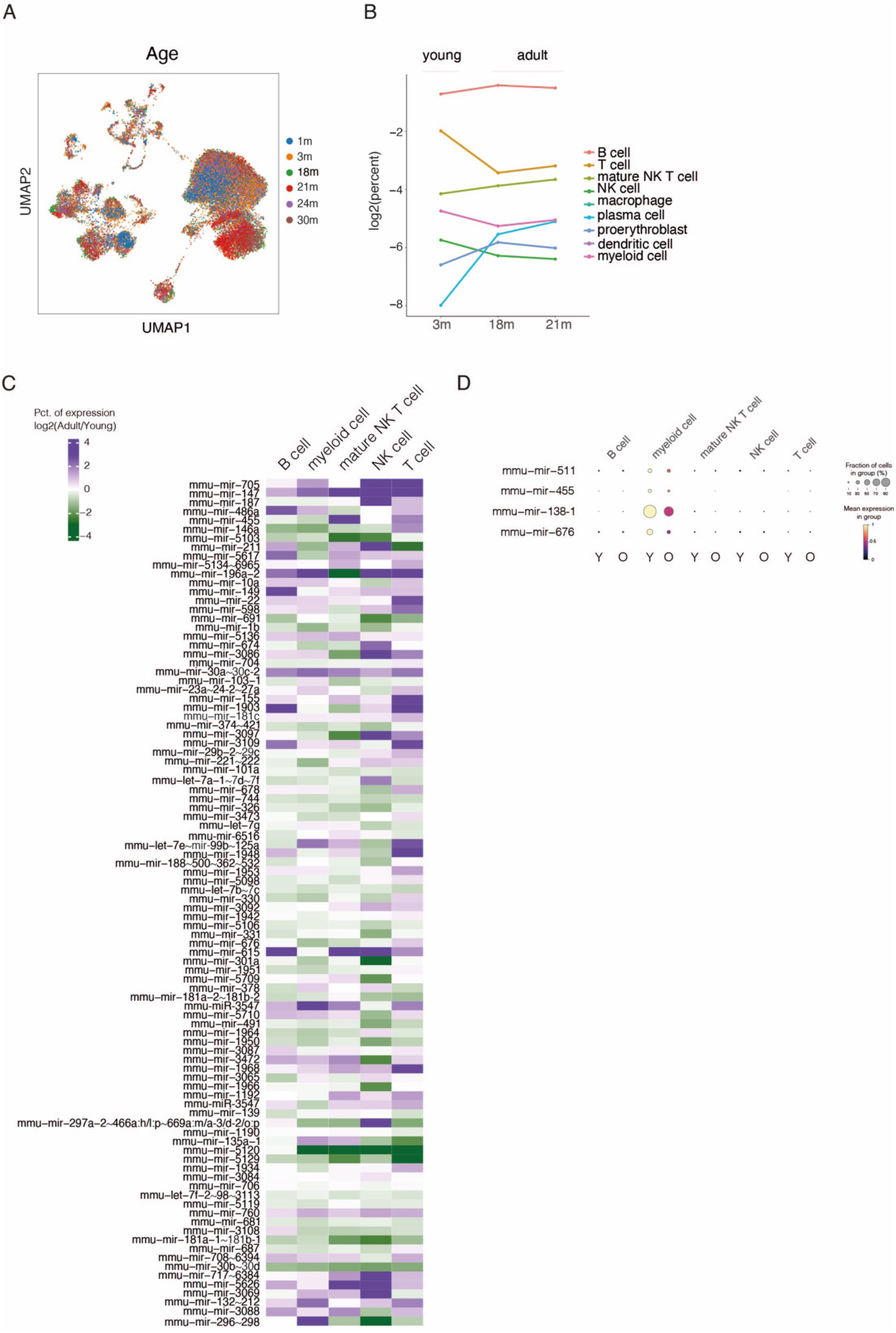
**Female analysis of splenic cell types in TMS data. (**A) UMAP of spleen tissue colored by age. (B) Cell type percentages from female mice present in each age group of the dataset are shown as a line plot. (C and D) Heatmap of miRNAs found concordantly regulated in splenic male samples during aging and plotted from female samples comprising young (3 months) and adult (18 and 21 months) mice.

**Figure S3.**
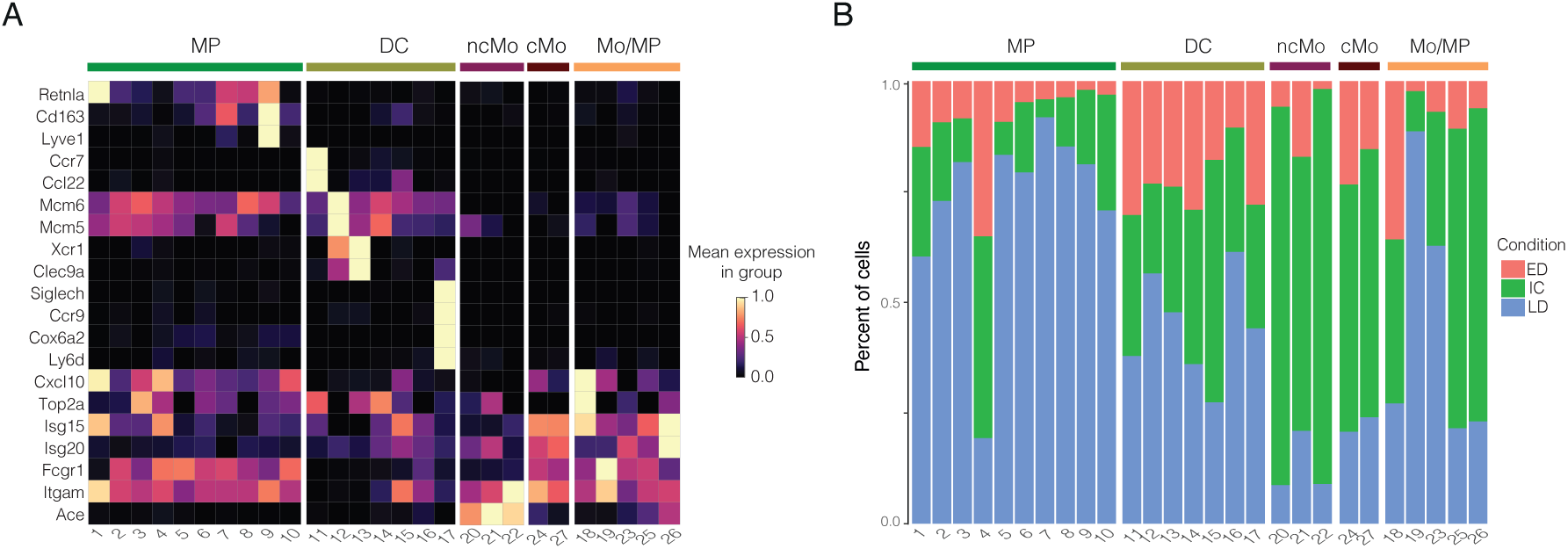
(A) Heatmap showing expression of literature-based gene markers in nodes. Expression is row-scaled. (B) Proportions of cells representing different conditions in each node.

**Figure S4.**
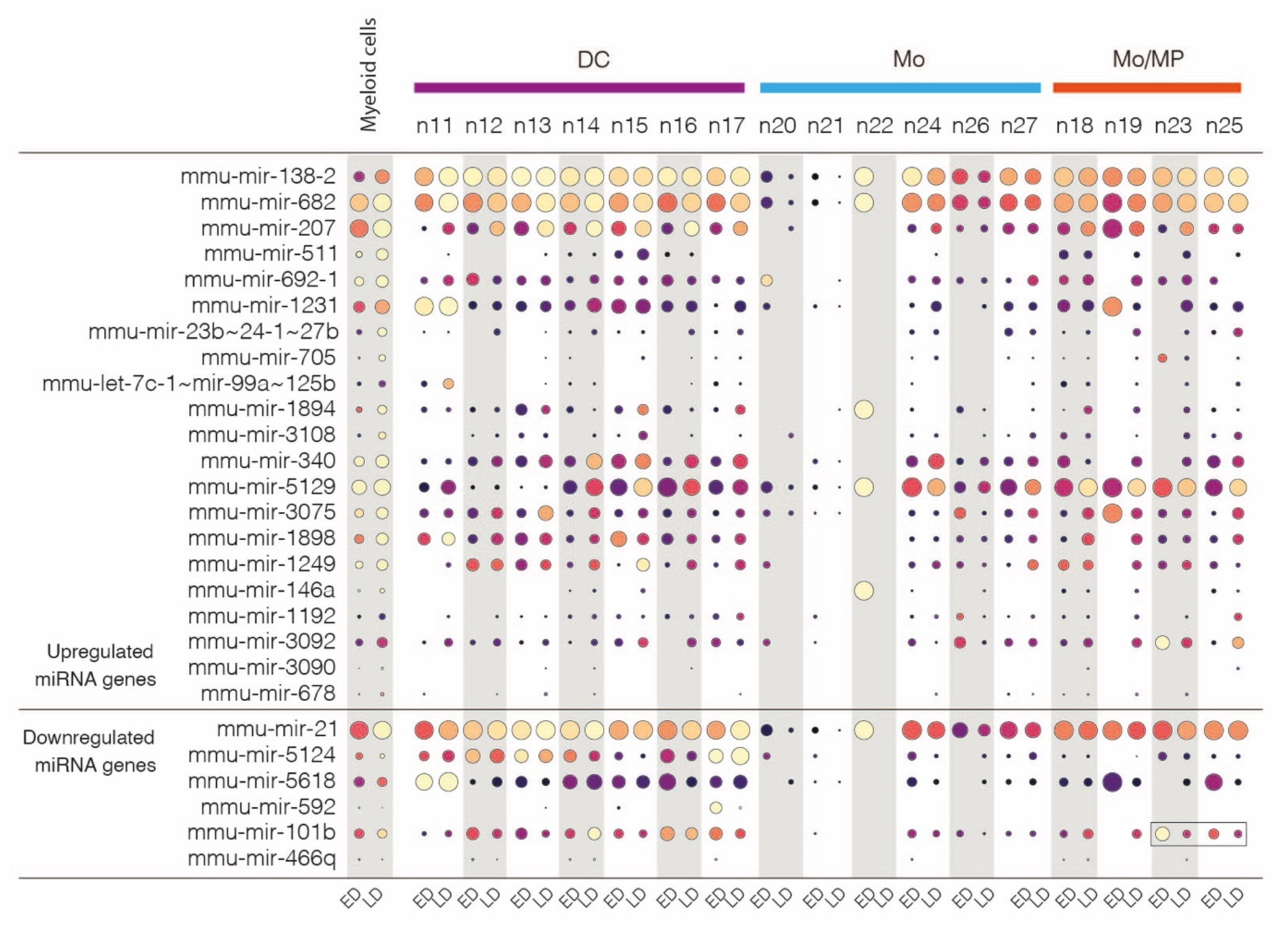
Changes in miRNA gene expression during disease progression in dendritic cells and monocytes. MiRNA genes with altered expression during disease progression are shown as in Fig. 4.

**Figure S5.**
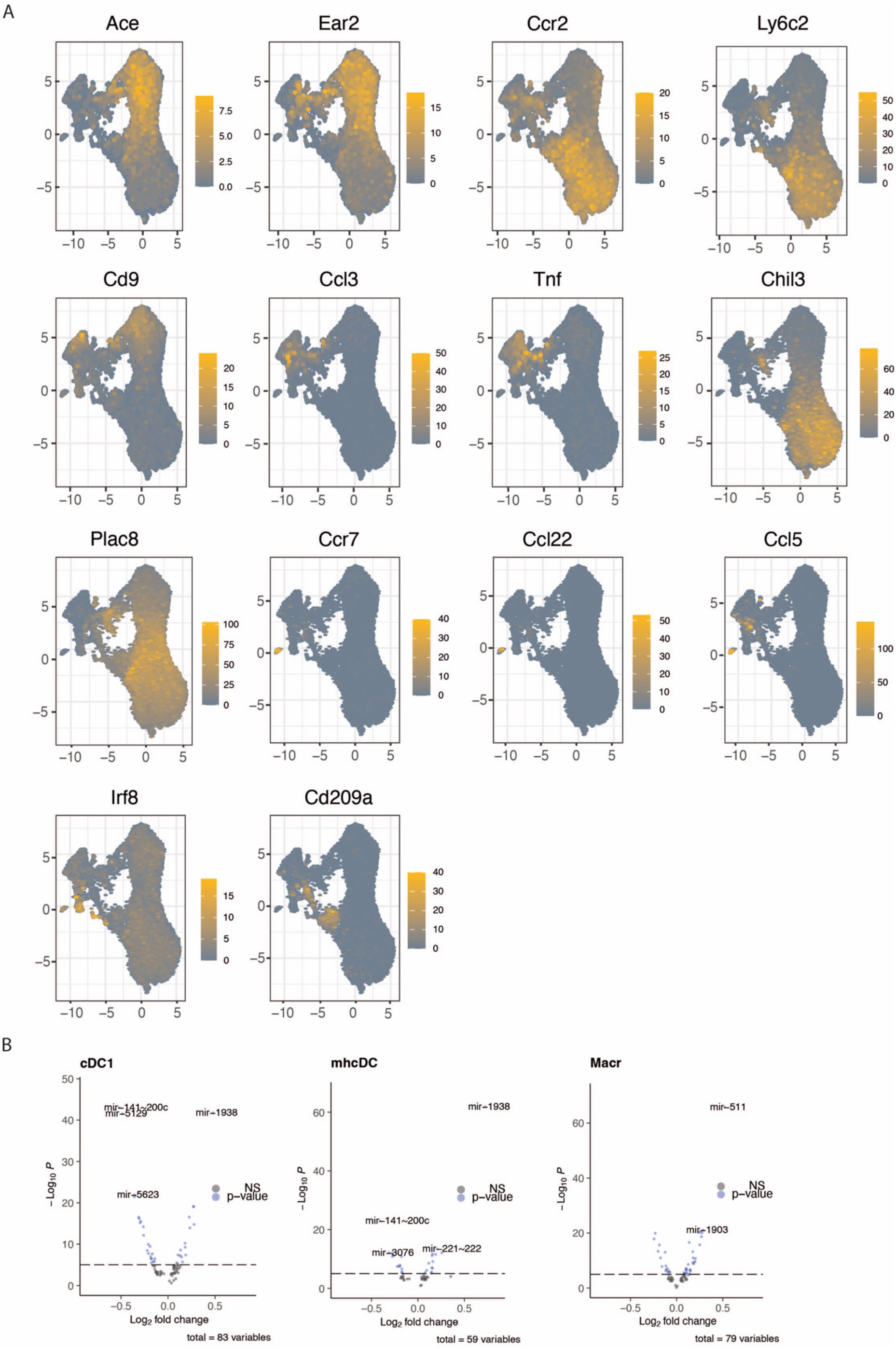
Marker genes for myeloid cell subpopulations. (A) Marker gene expression is shown on UMAP representing the tissue and blood myeloid cells (high expression in yellow tone). (B) Volcano plots representing significant miRNA gene loci from tissue vs blood comparison of monocyte subtypes. Top miRNA loci are indicated on the plot. FC correspond to differences in detection rate.

